# Model-based integration of citizen-science data from disparate sources increases the precision of bird population trends

**DOI:** 10.1101/2020.11.25.397380

**Authors:** Lionel R Hertzog, Claudia Frank, Sebastian Klimek, Norbert Röder, Hannah GS Böhner, Johannes Kamp

**Author notes:** **Correspondence**: Lionel R Hertzog. **Funding information**: Federal Ministry of Food and Agriculture (Germany).

## Abstract

**Aim:** Timely and accurate information on population trends is a prerequisite for effective biodiversity conservation. Structured biodiversity monitoring programs have been shown to track population trends reliably, but require large financial and time investment. The data assembled in a large and growing number of online databases are less structured and suffer from bias, but the number of observations is much higher compared to structured monitoring programs. Model-based integration of data from these disparate sources could capitalize on their respective strengths.

**Location:** Germany.

**Methods:** Abundance data for 26 farmland bird species were gathered from the standardized Common Breeding Bird Survey (CBBS) and three online databases that varied with regard to their degree of survey standardization. Population trends were estimated with a benchmark model that included only CBBS data, and five Bayesian hierarchical models integrating all data sources in different combinations. Across models, we compared consistency and precision of the predicted population trends, and the accuracy of the models. Bird species body mass, prevalence in the dataset and abundance were tested as potential predictors of the explored quantities.

**Results:** Consistency in predicted annual abundance indices was generally high especially when comparing the benchmark models to the integrated models without unstructured data. The accuracy of the estimated population changes was higher in the hierarchical models compared to the benchmark model but this was not related to data-integration. Precision of the predicted population trends increased as more data sources were integrated.

**Main conclusions:** Model-based integration of data from different sources can lead to improved precision of bird population trend estimates. This opens up new opportunities for conservation managers to identify declining populations earlier. Integrating data from online databases could substantially increase sample size and thus allowing to derive trends for currently not well-monitored species, especially at sub-national scales.

## Introduction

Biodiversity is undergoing rapid change worldwide due to anthropogenic pressures (Dornelas et al. 2013). Monitoring the trends of species populations across space and time is essential to assess the human impact on nature (Balmford et al. 2003), measure progress toward international biodiversity targets (Tittensor et al. 2014) and evaluate the effectiveness of policy interventions (Donald et al. 2007). Robust population monitoring is constrained by the availability of biodiversity data that varies considerably over space, time and with regard to the taxa covered (Amano et al. 2016). Long-term, systematic survey programs that employ a formal sampling design and a rigorous protocol often allow the estimation of accurate species population trends (Gregory et al. 2005, Boersch-Supan et al. 2019). Because these structured monitoring schemes rely on a large number of dedicated, skilled volunteers for fieldwork, they are resource-intensive, costly, and difficult to maintain in the long term (Schmeller et al. 2009). Structured data are available for only a small proportion of all taxa as monitoring schemes are usually restricted to comparatively wealthy and densely populated regions such as Europe and North America (Proença et al. 2017). Due to the limited number of sampling plots and often large variation in the data, the precision of the population trend estimates can be low (Snäll et al. 2011). As a consequence, weak population trends, especially of rare species, might be difficult to detect and the implementation of conservation actions delayed.

To meet obligations towards biodiversity conservation, governments, non-governmental organizations (NGOs) and researchers increasingly rely on less structured data, collected by large numbers of volunteers with varying levels of skill and submitted to public online citizen science databases (Sullivan et al. 2014, Amano et al. 2016, Chandler et al. 2017). These datasets have been either defined as “unstructured” when no or little ancillary data are collected, or as “semi-structured” when the design of the database allows to extract information on effort (e.g., route length, time spent surveying) or sampling completeness (e.g., when species “checklists” are collected that provide information on undetected species) (Sullivan et al. 2014, Kelling et al. 2019, Neate-Clegg et al. 2020). Public online databases that assemble biodiversity data have grown rapidly in recent years due to technological advances such as recording apps and audience-targeted advertising, e.g. via social media (Dickinson et al. 2012, Kelling et al. 2019). They provide larger amounts of data and greater spatial coverage compared to structured surveys, at lower cost and management effort (Sullivan et al. 2014, Amano et al. 2016, Chandler et al. 2017). While data feedback to the managing organization and the processing of structured data collected in large, coordinated monitoring schemes often takes one to two years until population trend estimates are available, biodiversity data in online databases is ready to use in real time, potentially allowing to obtain more timely estimates of population trends.

However, the usual absence of a survey design and the absence, or much less rigorous character of the sampling protocol in semi- and unstructured data leads to various biases such as uneven sampling over space and time (Isaac et al. 2014), uneven sampling effort per visit (Szabo et al. 2010) and varying detection and reporting probabilities (van Strien et al. 2013, Isaac & Pocock 2015). These biases, if not properly accounted for, can severely affect the reliability of population trends (Isaac et al. 2014). There is a growing set of advanced modelling approaches to account for some of the biases to provide reliable estimates of biodiversity trends (van Strien et al. 2013, Isaac et al. 2014). Recent examples include the development of ensemble models using spatio-temporal subsampling (Fink et al. 2020), or the use of occupancy models to simultaneously estimate trends for 31 taxonomic groups in the UK (Outhwaite et al. 2019).

Previous comparisons of trends from structured and unstructured datasets showed that negative trends estimated from structured monitoring were sometimes not picked up by unstructured data in the majority of species explored (Snäll et al. 2011, Kamp et al. 2016). This suggests that caution is needed when relying on online databases alone for trend estimation. However, good agreement was reached with “checklist” or semi-structured databases (where observers report all species), especially for common species (van Strien et al. 2013, Boersch-Supan et al. 2019).

Rather than analysing differently structured datasets separately, joint modelling or data integration approaches have recently been advanced as an effective means to use these disparate data in a coherent manner (Miller et al. 2019, Isaac et al. 2020). These approaches combine the strength of structured datasets (e.g. little locational bias) with those of semi- and unstructured datasets (e.g. large number of records and broad coverage). By building models relying on a substantially increased amount of data, the precision of population trend estimates may also increase, leading to less uncertainty around the trend estimates. Therefore, negative population trends might be identified earlier than with structured monitoring alone (Borsch-Supan et al. 2019). Moreover, the larger spatial coverage provided by semi- and unstructured datasets may increase the level of agreement between the predicted and observed values leading to higher accuracy of the trends. Finally, species with low to medium abundance and prevalence (i.e. proportion of sampling sites were the species was observed), often of conservation concern, are poorly covered in structured large-scale monitoring schemes. The estimation of their population trend is therefore difficult (Snäll et al. 2011). Joint models could potentially allow the derivation of robust trends for these species by efficiently using all available data.

Recently developed data integration models explicitly consider that the process of interest (i.e. temporal changes in bird populations) is being measured through different sampling regimes (Miller et al. 2019, Isaac et al. 2020). Through larger spatial coverage and explicit modelling of different sampling processes, model-based data integration results in greater accuracy and precision of the derived quantities (Bowler et al. 2019, Robinson et al. 2020). However, data integrated models still need to be tested across a broader range of taxa and at wider spatial scales to assess the potential advantages of the approach over more established modelling frameworks (Isaac et al. 2020, Simmonds et al. 2020).

Here, we apply model-based data integration combining data from a structured monitoring program with semi- and unstructured data from several online databases. We used data on birds, one of the best monitored taxonomic groups that is often used as indicator of environmental change (Donald et al. 2007, Jørgensen et al. 2016). To reveal long-term trends in bird populations, we used high-quality, structured citizen science data from the Common Breeding Bird Survey (CBBS) in Germany. This long-term monitoring scheme is based on data collected annually on up to 1700 stratified randomly selected plots, with standardized survey effort (Kamp et al. 2020). Next to this structured dataset, data from three online databases were extracted, the Germany-wide *ornitho.de*, and the global *ebird.org* and *observation.org*. These databases contain large quantities of bird records collected without sampling design and with no (i.e. unstructured) or light (i.e. semi-structured) sampling protocol. To assess the usefulness of data integration for estimating bird population trends, we tested different data integration scenarios ranging from no integration at all to all datasets being integrated in one common model. We compared the consistency of model outcomes such as annual abundance indices and temporal trends to those of simple yet well-established log-linear models for structured count data.

We predict that (a) model-based integration of citizen-science data from databases with varying degree of structuredness generally improves precision and accuracy of population trends, and (b) the gains in precision and accuracy vary across species and are explained by species characteristics such as prevalence, abundance and body mass, with greater improvements for rarer species.

## Methods

### Structured monitoring data

Structured, plot-specific data from the German Common Breeding Bird Survey (CBBS) were available for the years 2005 to 2018. The CBBS follows a rigorous protocol in which experienced volunteers record breeding birds along a fixed route of approximately 3 km length within a 1 km^2^ sampling plot on four annual visits during the breeding season (10 March to 20 June, Kamp et al. 2020). An abundance estimate per species and plot is derived by combining the data of the four survey rounds into “territories” following Bibby (2000). The sampling plots were selected randomly stratified, with strata mirroring Germany’s environmental regions and habitats (Kamp et al. 2020). For more detail on the survey design and sample sizes see Kamp et al. (2020) and Appendix Text S1. From the CBBS, we selected 26 farmland birds, i.e. species associated with farmland as breeding and or feeding habitat (Table 1). This species group suffered the strongest population declines in Europe during the twentieth century, and is hence in the focus of conservation in Germany (Busch et al. 2020, Kamp et al. 2020) and beyond (Gregory et al. 2019). Farmland birds also exhibit higher functional and phylogenetic diversity than forest bird assemblages in Europe (García-Navas et al. 2020) and a high degree of specialization (Kirk et al. 2020), and should thus respond in an especially sensitive way to environmental change.

**Table 1:**
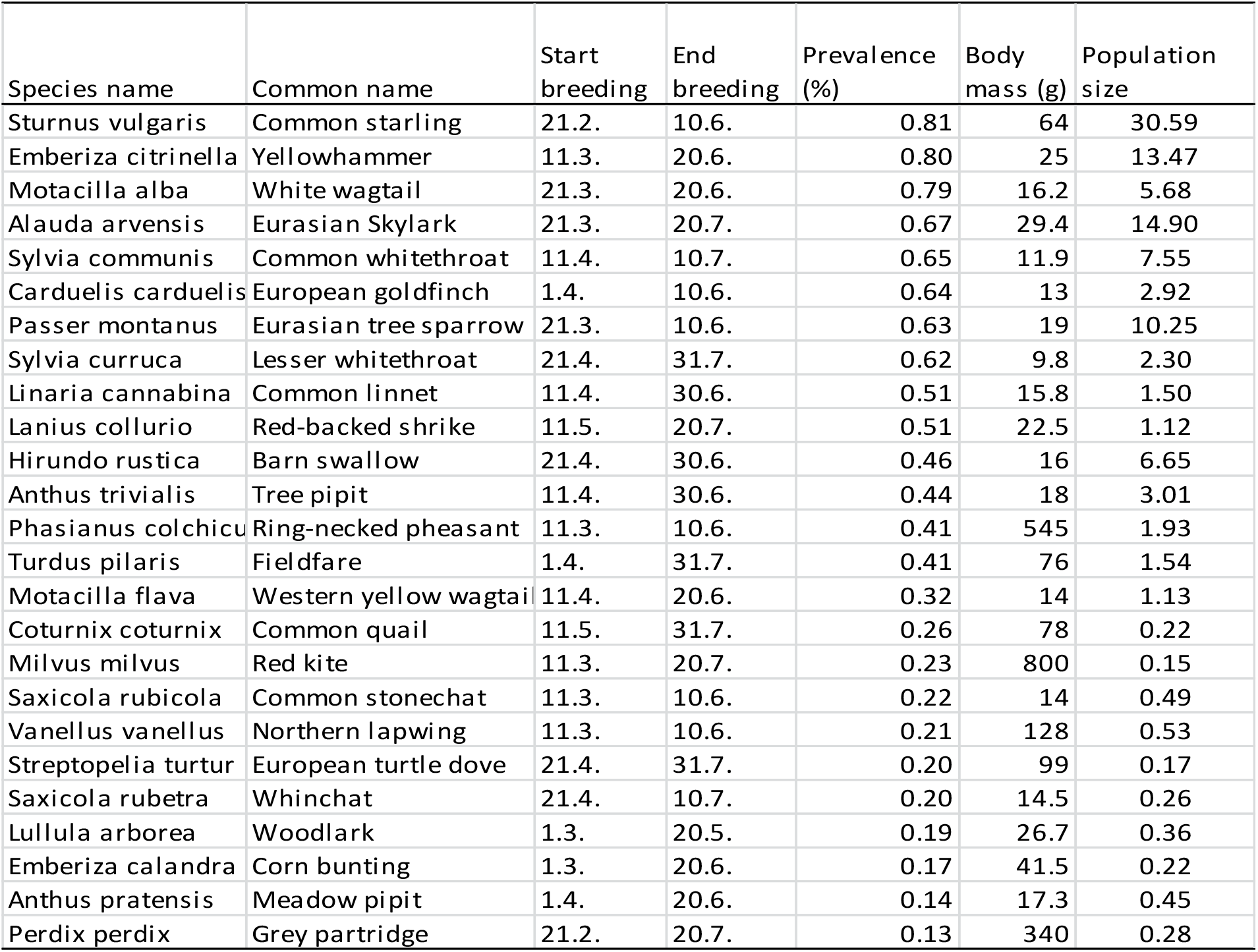
List of the 26 selected farmland bird species with information on start and end of the breeding period, ordered by prevalence (computed as the proportion of CBBS routes where the species was recorded). Mean body mass and estimated national population size (number of breeding birds in 100’000) are also given.

| Species name | Common name | Start breeding | End breeding | Prevalence (%) | Body mass (g) | Population size |
| --- | --- | --- | --- | --- | --- | --- |
| *Sturnus vulgaris* | Common starling | 21.2. | 10.6. | 0.81 | 64 | 30.59 |
| *Emberiza citrinella* | Yellowhammer | 11.3. | 20.6. | 0.80 | 25 | 13.47 |
| *Motacilla alba* | White wagtail | 21.3. | 20.6. | 0.79 | 16.2 | 5.68 |
| *Alauda arvensis* | Eurasian Skylark | 21.3. | 20.7. | 0.67 | 29.4 | 14.90 |
| *Sylvia communis* | Common whitethroat | 11.4. | 10.7. | 0.65 | 11.9 | 7.55 |
| *Carduelis carduelis* | European goldfinch | 1.4. | 10.6. | 0.64 | 13 | 2.92 |
| *Passer montanus* | Eurasian tree sparrow | 21.3. | 10.6. | 0.63 | 19 | 10.25 |
| *Sylvia curruca* | Lesser whitethroat | 21.4. | 31.7. | 0.62 | 9.8 | 2.30 |
| *Linaria cannabina* | Common linnet | 11.4. | 30.6. | 0.51 | 15.8 | 1.50 |
| *Lanius collurio* | Red-backed shrike | 11.5. | 20.7. | 0.51 | 22.5 | 1.12 |
| *Hirundo rustica* | Barn swallow | 21.4. | 30.6. | 0.46 | 16 | 6.65 |
| *Anthus trivialis* | Tree pipit | 11.4. | 30.6. | 0.44 | 18 | 3.01 |
| *Phasianus colchicus* | Ring-necked pheasant | 11.3. | 10.6. | 0.41 | 545 | 1.93 |
| *Turdus pilaris* | Fieldfare | 1.4. | 31.7. | 0.41 | 76 | 1.54 |
| *Motacilla flava* | Western yellow wagtail | 11.4. | 20.6. | 0.32 | 14 | 1.13 |
| *Coturnix coturnix* | Common quail | 11.5. | 31.7. | 0.26 | 78 | 0.22 |
| *Milvus milvus* | Red kite | 11.3. | 20.7. | 0.23 | 800 | 0.15 |
| *Saxicola rubicola* | Common stonechat | 11.3. | 10.6. | 0.22 | 14 | 0.49 |
| *Vanellus vanellus* | Northern lapwing | 11.3. | 10.6. | 0.21 | 128 | 0.53 |
| *Streptopelia turtur* | European turtle dove | 21.4. | 31.7. | 0.20 | 99 | 0.17 |
| *Saxicola rubetra* | Whinchat | 21.4. | 10.7. | 0.20 | 14.5 | 0.26 |
| *Lullula arborea* | Woodlark | 1.3. | 20.5. | 0.19 | 26.7 | 0.36 |
| *Emberiza calandra* | Corn bunting | 1.3. | 20.6. | 0.17 | 41.5 | 0.22 |
| *Anthus pratensis* | Meadow pipit | 1.4. | 20.6. | 0.14 | 17.3 | 0.45 |
| *Perdix perdix* | Grey partridge | 21.2. | 20.7. | 0.13 | 340 | 0.28 |

### Semi-structured and unstructured data from online databases

For the selected farmland birds, we harnessed records from a number of public online citizen science databases. In contrast to structured monitoring data, these databases are not based upon a systematic site-selection and lack a rigorous sampling protocol. Two broad types of data are available from online databases: (i) semi-structured and (ii) unstructured. Semi-structured data are characterized by the availability of some ancillary data on the observation process that allows to quantify effort and bias. Unstructured data represent incidental records without measures of effort, or information on species not observed (Kelling et al. 2019).

The database ornitho (www.ornitho.de) is a Germany-wide platform allowing observers to report bird observations across Germany. On 08/10/2020, the database contained 50.9 million records entered by *ca*. 21,200 active observers. Observers can choose to enter observations as incidental records (unstructured element of the database), or enter a complete checklist of observations (semi-structured element). Incidental records and complete checklists are assigned precise coordinates, or are allocated to the centre of a grid cell of approximately 1 x 1 km (following the German “Halbminuten”-grid). Additional information on the start and end time need to be provided in semi-structured data and time spent observing birds can thus be used as a measure of effort.

eBird (www.ebird.org) is the largest global database of bird observations (Sullivan et al. 2014, Neate-Clegg et al. 2020), with >500 million records on 07/10/2020. Observations can be filtered to separate unstructured (incidental sightings) from semi-structured (complete checklists) data. Observers on eBird are required to assign their data to four categories: traveling (walked route of preferentially < 5 miles, with information on duration), stationary (watching from a fixed location), historical (retrospective data entry) and incidental (birdwatching was not the primary purpose). The first two categories are semi-structured data while the latter two are unstructured data. For the traveling and stationary checklists, effort information (time, duration, party size and distance travelled) is requested from the recorder.

The Netherlands-based platform observation (observation.org) is global in scope and allows the entry of data on 19 taxonomic groups (https://observation.org/statistiek.php), with birds, plants, butterflies, mammals and dragonflies contributing the bulk of the data. There were around 40 million records on 07/10/2020 (https://www.gbif.org/publisher/c8d737e0-2ff8-42e8-b8fc-6b805d26fc5f), most of which were from the Netherlands, with 70% bird observations. The data are unstructured, at least for birds, since the reporting of effort measures is not mandatory, and no checklist option is available.

Records from semi- and unstructured records in eBird and observation were mapped onto the German “Halbminuten”-grid used in ornitho.

### Data pre-processing

To increase comparability across the different datasets, pre-processing was applied for the online databases (ornitho, eBird and observation). We included only data from the years 2012–2019 because only 3.6% of the observations were older than 2012, for all databases combined (see also Fig. S1). As birds on migration could lead to distortions in breeding bird abundance, we considered only those observations that fell into species-specific breeding times by filtering out records outside of a defined breeding period, following Südbeck et al. (2005) (Table 1). Non-stationary birds (i.e. flyovers) were also removed from the unstructured data from ornitho. Records from online databases where the reported bird abundance was larger than the annual maximum observed for that species in the CBBS were also dropped to prevent the inclusion of clear outliers such as flocks of 10,000 starlings during the breeding season. Records from all German islands were removed to reduce locational bias that might have been introduced as these islands are hotspots of birdwatching attracting a large number of people, and as they are particular dynamic habitats. Where several observations per locality (grid cell), species and year were available, only the maximum number of simultaneously observed individuals during breeding season was retained. This site maximum is a good approximation of bird abundance at the locality. To correct for varying sampling effort in semi-structured data (complete lists from ornitho and eBird), we used the duration of the observation period in hours as a measure of effort (Szabo et al. 2010). For all unstructured data (incidental sightings from ornitho, eBird and observation), we used the total number of farmland bird species recorded on that day, at that locality by that observer as a proxy for the sampling effort. After these pre-processing steps, the datasets consisted of 1699 routes from the CBBS, 23,194 and 5600 sites respectively from the ornitho and eBird semi-structured data and 183,057, 2134 and 7954 sites respectively from ornitho, eBird and observation unstructured data (Fig. 1).

**Figure 1:**
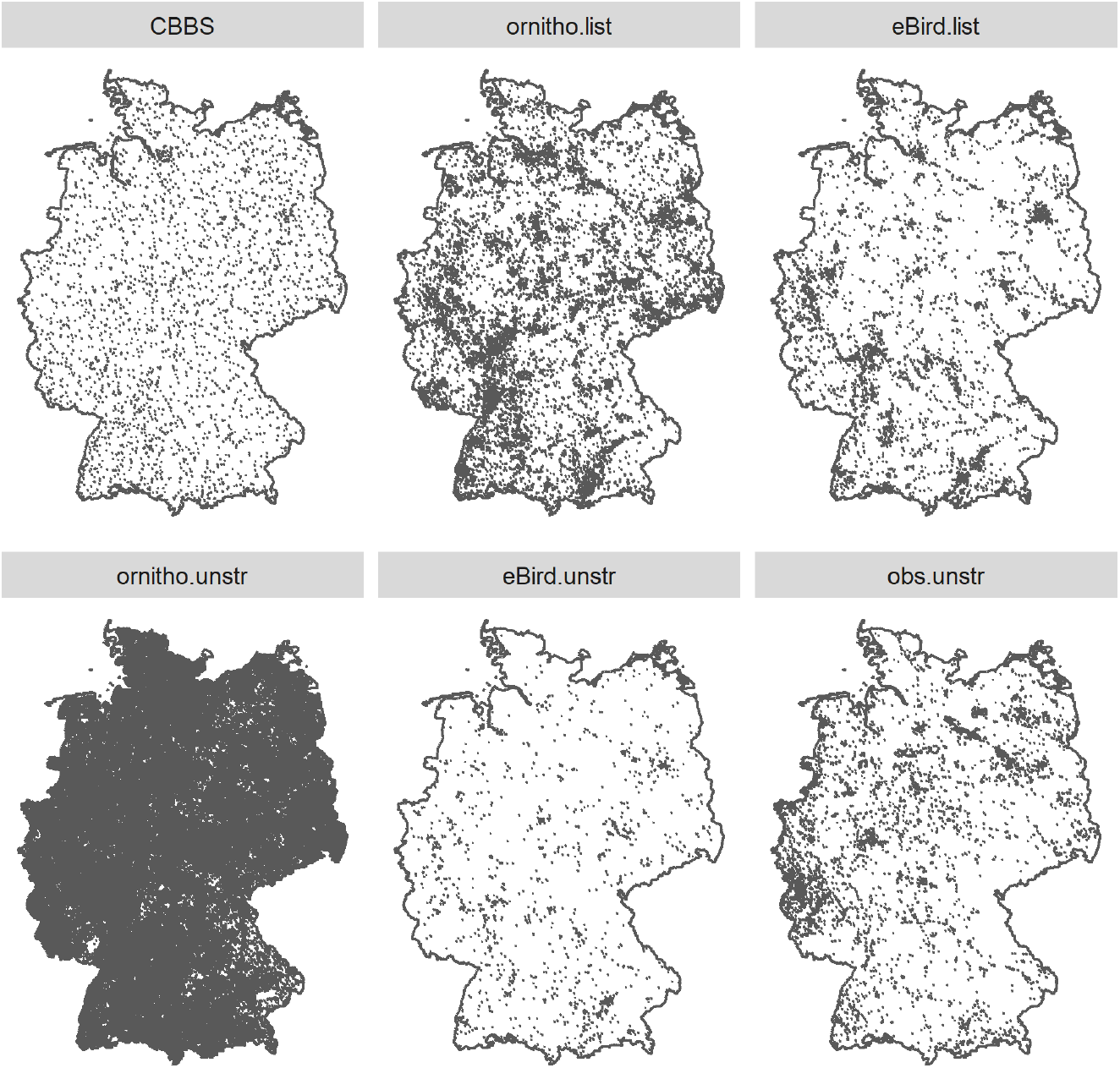
Spatial distribution of the sites used in the analysis from the various datasets. CBBS: plots monitored through the Common Breeding Bird Survey, ornitho.list: complete checklists from the ornitho database, eBird.list: complete checklists from the eBird database, ornitho.unstr: unstructured records from ornitho, eBird.unstr: unstructured observations from eBird, obs.unstr: unstructured observations from observation.

### Modelling approach

Six models were fitted to data of all 26 species separately to evaluate the effect of different data integration strategies on consistency and precision of the predicted population trends, and on model accuracy. In all models the response variable was the yearly bird abundance either for the CBBS routes or for the grid cells. Two broad class of models were used: a log-linear Poisson regression method (“TRIM”, see below) and five different Bayesian hierarchical generalized linear models (Model 2-6).

#### Model 1 - TRIM

TRIM (TRends and Indices for Monitoring data) is a software developed in 1991 to analyse temporal variation in biodiversity monitoring data (van Strien et al. 2004). TRIM implements a loglinear Poisson regression and Generalized Estimating Equations to produce yearly abundance indices and trends, and is a standard tool employed to estimate bird population trends across most European countries. The algorithm is now available as an R package (rtrim, Bogaart et al. 2018). Due to its widespread use across Europe, we consider this model as the “benchmark” in our comparisons. Various model variants are available in rtrim and we used model 3 on the CBBS data which models the observed counts as a function of site-varying and year-varying effect:

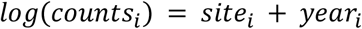

An overdispersion term was generally included for all species as overdispersion is a common problem when fitting models to count data with a Poisson error term. For some species the dispersion parameter was estimated to be below or around 1. These models were then refitted without the correction for overdispersion. From the TRIM models we derived (a) the R-square computed following (Effron 1978), (b) the normalized root mean squared error (the root mean squared error divided by the average count) and (c) the predicted total number of birds per year, so the summed breeding bird abundance across CBBS routes for each year, together with the 95% confidence bands.

#### Models 2-6 - Hierarchical generalized linear models (with data integration)

Models 2 to 6 were fitted using the R package ‘*greta*’ (Golding 2019) which allows flexible model definition and parameter estimation using a Bayesian approach with a Hamiltonian Monte Carlo algorithm. For all of these models the response variable was the bird abundance. The explanatory variables were the year of the observation as a fixed effect (categorical, as in Model 1), and the location of the observation (CBBS route or grid cell ID) as a random intercept. For the observations based on complete lists (semi-structured data) from eBird or ornitho an offset term was added to account for the variation in sampling effort (see above). In these models, observations were modelled relative to the effort that went into the data recording. For the unstructured data from eBird, ornitho and observation the number of species recorded during the observation event was included as a covariate to account for varying sampling effort, site suitability and observer expertise. Different datasets were integrated in these models (see Table 2 for an overview of which dataset were integrated in the different models). A full description of further model specifications, such as prior definitions and sampling settings, is given in Appendix Text S2.

**Table 2:** Overview of the structure of the different datasets and their use in the different models. Model 1 was fitted with the rtrim R-package while models 2-6 were fitted with the greta R-package.

| Dataset | CBBS | ornitho |  | eBird |  | observation |
| --- | --- | --- | --- | --- | --- | --- |
| Dataset type | structured | Semi-structured<br>(complete checklists) | unstructured | Semi-structured<br>(complete checklists) | unstructured | unstructured |
| Model 1 (TRIM) | x |  |  |  |  |  |
| Model 2 | x |  |  |  |  |  |
| Model 3 | x | x |  |  |  |  |
| Model 4 | x | x | x |  |  |  |
| Model 5 | x | x |  | x |  |  |
| Model 6 | x | x | x | x | x | x |

### Assessing trend consistency

Based on the model estimated parameters, the number of birds was predicted for each CBBS route in each year between 2005 and 2018 in model 1 (TRIM model) and model 2 (greta CBBS model). For the other models, the predicted yearly variations were estimated between 2005 and 2019 for all CBBS routes but also for the grid cells with observations. To be consistent across the models only predictions on the CBBS routes were used. The yearly predicted bird numbers were summed across the CBBS routes per year yielding an abundance index, with an associated 95% confidence band. To check for consistency in the predicted trends across the different models two approaches were used: First, trends were classified as follows: if the abundance index in 2018 was below the lower confidence limit for the abundance index in 2012, the trend was classified as decreasing, if the 2018 prediction was above the upper confidence limit for 2012, the trend was classified as increasing, otherwise the trend was classified as stable. The classification of the trends was then compared between model 1 (TRIM) and the other models across the 26 species to identify potential discrepancies. Second, for the years between 2012 and 2018 we calculated the Spearman rank correlation coefficients between the yearly abundance indices from model 1 and from the other models for the respective year and species. Correlation coefficients above 0.7 were assumed to represent consistency in the derived abundance indices and their temporal fluctuations. The correlation coefficients were further plotted against the three selected species characteristics (prevalence, abundance and body mass, see below) to explore whether these characteristics explained variation in the across-model consistency.

### Assessing accuracy and precision

The accuracy of the model to predict the breeding bird number on the CBBS routes was estimated by computing the normalized root mean squared error as follows (higher values meaning less accurate models):

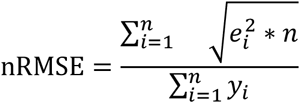

where *e* are the model residuals (considering only the CBBS routes) and *y* are the observed values.

The precision of the model predictions was derived by averaging the confidence ranges around the abundance indices between 2012 and 2018. These values were normalized with the average species-specific mean abundance index in order to be comparable across species. The accuracy and precision values of models 2 to 6 were compared to those of the model 1 by using percent changes. To investigate whether the consistency, the accuracy and the precision of the predicted population trends was dependent on species characteristics (Table 1), we gathered information on species body mass (from Wahl et al. unpubl.) that can be used as a proxy for detectability (Johnston et al. 2014), mean national population estimates as a proxy for species abundance (Gerlach et al. 2019) and calculated prevalence (the proportion of CBBS routes were the species was observed).

## Results

### General model fitting

Good convergence values (Rhat < 1.1) were reached for all model parameters and for all species for models 2 to 5. Model 6 (integrating all datasets) showed poor convergence across multiple parameters for most of the species. Results from these models are therefore not reported. The abundance indices for each species between 2005 and 2019 are available in Appendix Figures S13 to S39.

### Trend consistency

The trend classification showed perfect agreement when comparing the classification from Model 1 (TRIM) to the hierarchical model only with CBBS data (Model 2, Fig. 2a). For Models 3 and 5 (CBBS + semi-structured data from ornitho and ornithon+ebird respectively), more than 75% of the species were classified similarly when compared to Model 1 (Fig. 2b-d). For four species that were classified as having stable trends under Model 1, Model 3 and 5 predicted increase or decrease in populations. For these two models only one species, Blackbird, showed inconsistent results where Model 1 predicted population decrease while Model 3 and 5 predicted population increases between 2012 and 2018. For Model 4 (CBBS + structured & unstructured ornitho data), only half of the species were classified similarly when compared to Model 1 (Fig. 2c). Four species showed trend classification in opposite directions, these were Common whitethroat, Barn swallow, Blackbird and Common stonechat.

**Figure 2:**
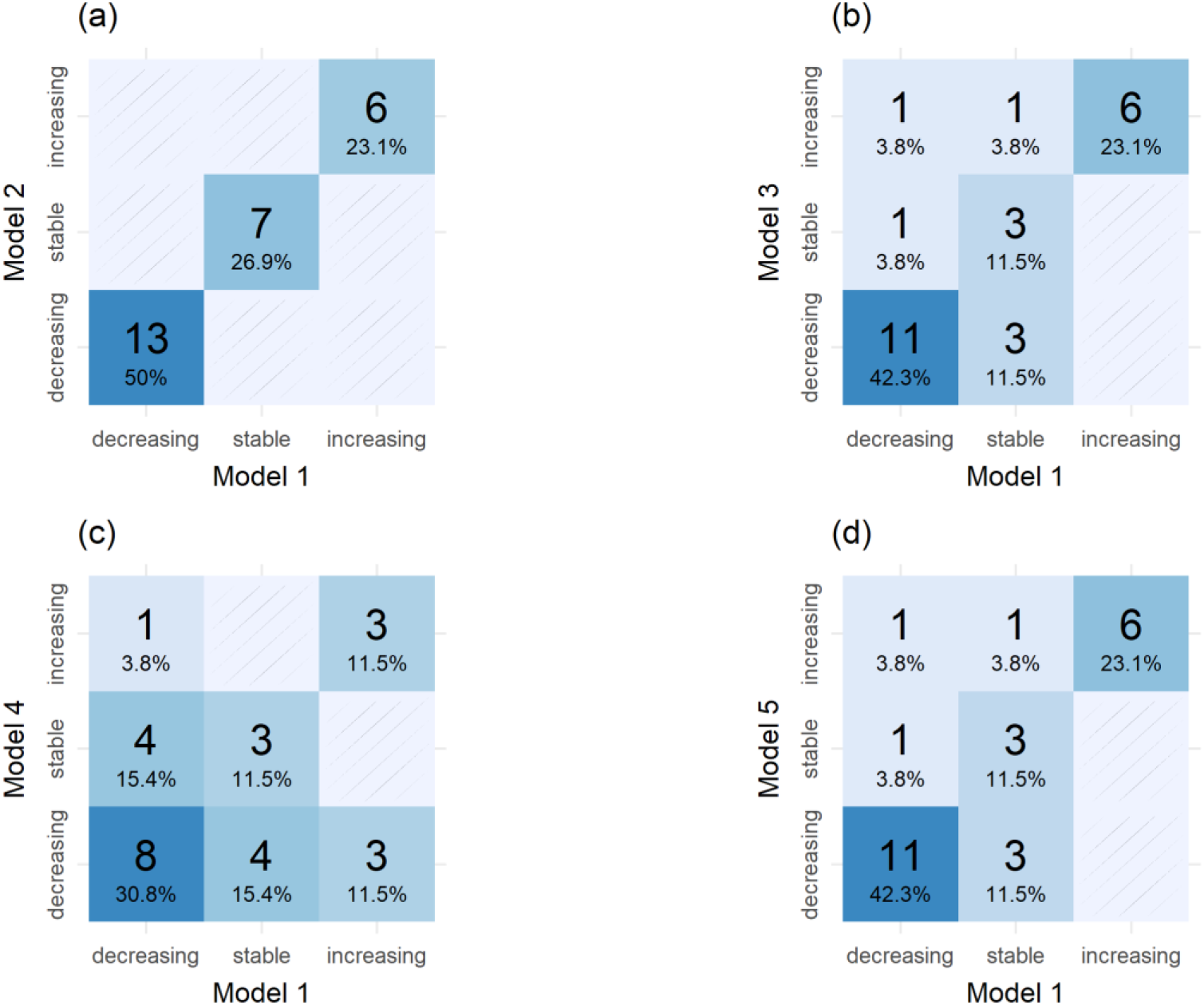
Confusion matrices of the trend classification, comparing Model 1 with Models 2-5 alternatingly. The percentage and numbers represent the proportion and the number of species falling in the different cases.

The annual abundance indices over the period 2012 to 2018 from model 1 were strongly correlated to trends from model 2 (mean Spearman rank correlation coefficient: 0.98, range 0.89 – 1.00, n = 26, Fig. 3), suggesting that the patterns in bird trends revealed by the standard TRIM approach can also be retrieved with models that differ in structure and fitting process. Model 3 and 5 that integrated the semi-structured data with the CBBS data were highly correlated with the predicted annual abundance indices from model 1 (15 and 13 out of the 26 bird species showed correlations above 0.7, respectively). For model 4 that integrated both, semi-structured and unstructured data, only 5 out of the 26 species showed good agreement in the predicted annual abundance indices between the CBBS model and the data integrated model. Overall, five species showed good consistency (Spearman correlation coefficient > 0.7) in the predicted abundance indices across the different models, these were: Common Quail, Turtle Dove, Ortolan bunting, Yellowhammer and Skylark. Eleven species showed a poor correlation (spearman correlation coefficient < 0.7) between model 1 and the data integrated models 3-5, these species were: White Wagtail, Whinchat, Tree Sparrow, Woodlark, Northern Lapwing, Barn Swallow, Grey Partridge, Red Kite, Starling, Meadow Pipit and Yellow Wagtail. The correlation between the CBBS and the data integrated models did not depend on the prevalence or the national population estimates of the species nor on the species body mass (Fig. S2 to S4).

**Figure 3:**
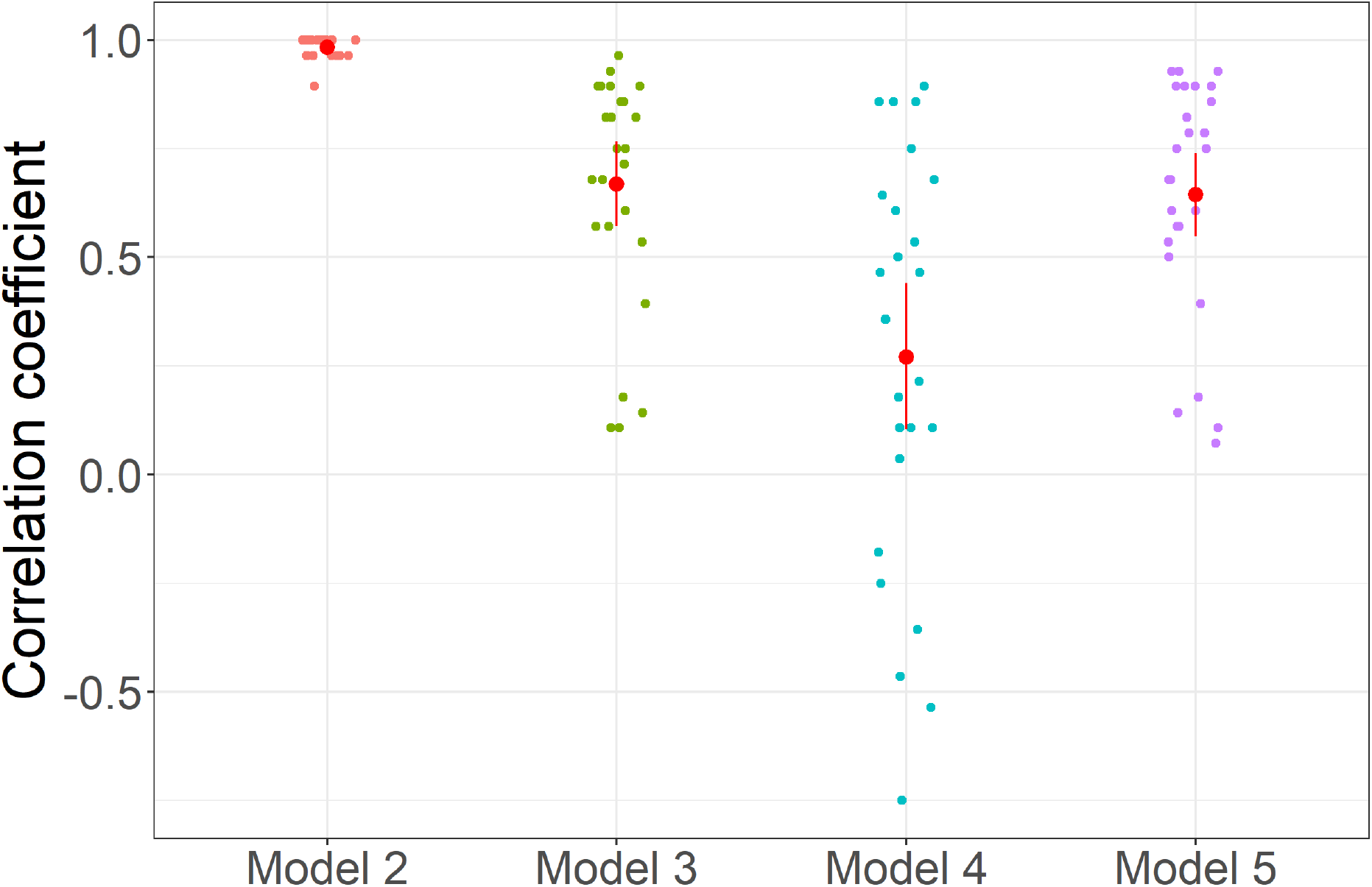
Correlation of predicted population annual abundance indices from the Bayesian Hierarchical Models 2-5 with those from Model 1 as measured by the Spearman correlation coefficient. The dots represent the selected bird species (n=26) and the red dots represent the mean values together with a bootstrapped 95% confidence interval. For the correlation only the years between 2012 and 2018, when data integration affected the predicted trends, are considered.

### Accuracy and precision

The Bayesian hierarchical models (model 2-5) tended to provide more accurate predictions than model 1 by an average of 25% and this was irrespective of the use of data integration (Fig. 4). The increase in accuracy in the hierarchical models compared to TRIM depended on the prevalence and the national population estimates of the species in the CBBS routes. The least abundant species showed an increase of 60% in accuracy in the hierarchical models compared to TRIM, while for the most common species the hierarchical models only led to a marginal increase in accuracy (Fig. S5-S6). Species body mass did not affect accuracy differences (Fig. S7).

**Figure 4:**
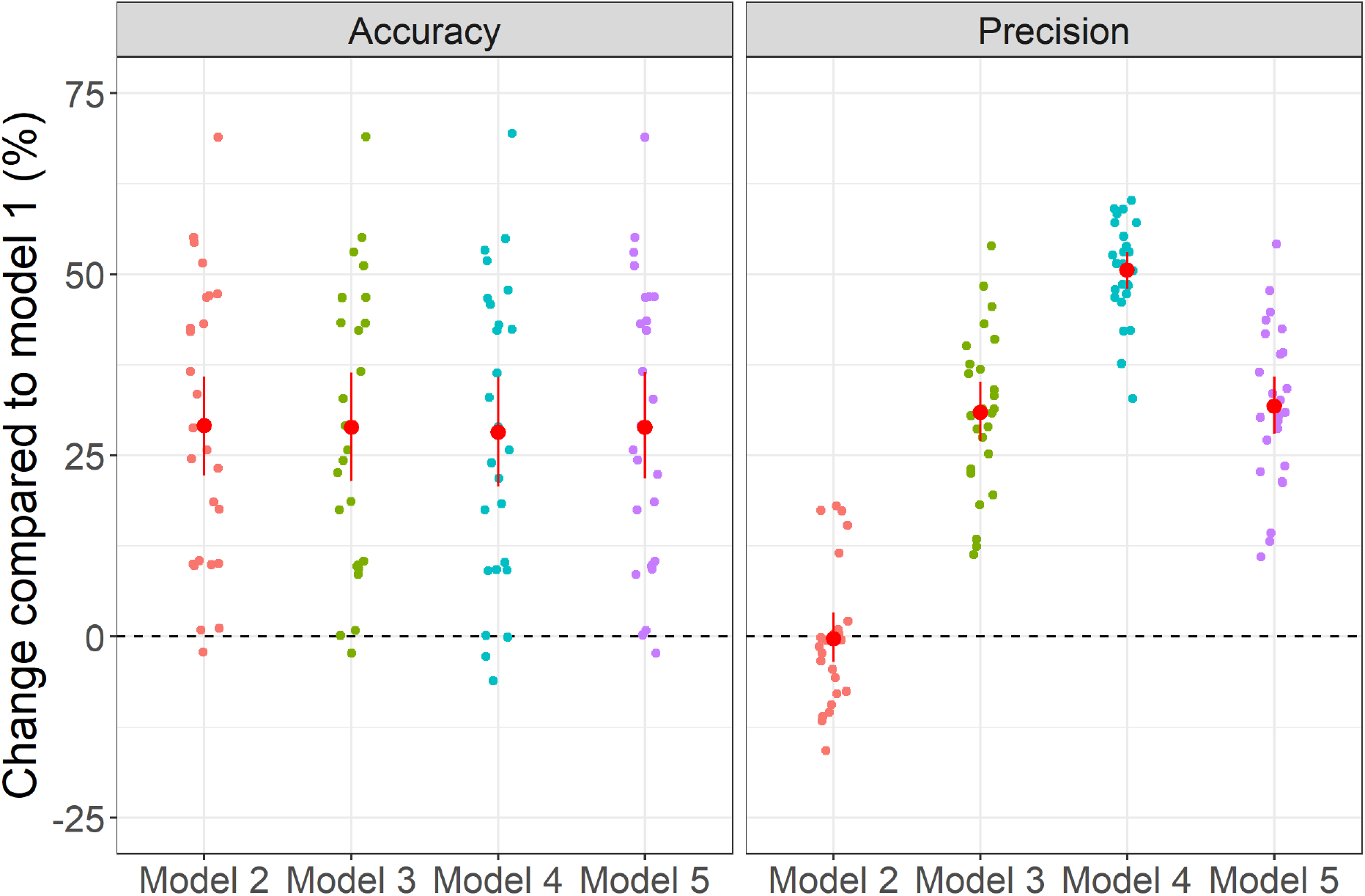
Effect on data integration on accuracy (left) and precision (right) of the predicted population trends measured as the percent changes between the Bayesian Hierarchical Models and model 1. Each dot represents one bird species (n=26), the red dots are the mean together with a 95% bootstrapped confidence interval.

Data integration increased the precision of the predicted annual abundance indices (Fig. 4). Adding the semi-structured data to the CBBS data increased the precision by around 25% compared to model 1. The most complex models integrating CBBS, semi-structured and unstructured data (model 4) increased the precision by around 50% compared to model 1. The national population estimates were in general not correlated with the changes in precision, only in the model integrating CBBS, semi-structured and unstructured data (Model 4) a negative correlation between abundance and precision changes was apparent (Fig. S8). On the other hand, the increase in precision in the data integrated models was not found to be related to species prevalence in the CBBS routes or to species body mass (Fig. S9, S10).

## Discussion

Timely and accurate information on population trends is of utmost importance to conservation managers and policy-makers to facilitate the development and implementation of effective conservation actions (Balmford et al. 2003, Chandler et al. 2017). Here, we investigated in which ways the integration of semi-structured data (checklists) and unstructured data from several online databases with structured data from highly standardized monitoring schemes affected temporal trends of 26 farmland bird species. Data integration increased the precision but not the accuracy of temporal trends. The use of Bayesian hierarchical models increased the accuracy of the temporal trends overall and especially for the rarer species. This was irrespective of the use of data integration. Integrating semi- and unstructured data from online databases with structured data led to discrepancies in the predicted fluctuations in annual abundance indices in at least 11 out of the 26 focal species, but when focusing on trend classification rather than on year-to-year variation good agreement was found between the different models especially for the models not integrating unstructured data.

### Different trends with different data integration

Comparing the different models revealed low correlations in annual abundance indices for 11 species between the models including only data from structured monitoring and the models integrating semi- and unstructured data. Disparities in trends could result from locational bias that varies with data source. Sampling locations are pre-selected in structured monitoring schemes, whereas participants in online databases choose freely where to observe birds (Keeling et al. 2019). Bias could result from changes in observer patterns and resulting locational bias over time (Boersch-Supan unpubl.). An increase of records near urban areas in recent years might have been caused by a greater involvement of occasional (as opposed to semi-professional) birdwatchers, triggered by the availability of recording apps. Because these areas are less likely to hold large numbers of the farmland birds under study here, temporal shifts in sampling locations with an increasing amount of abundance data from non-farmland sites could negatively bias the estimates from the structured monitoring in the integrated models. Graphical checks of the human population density distribution at the surveyed CBBS routes and from the online databases (see Text S.3) revealed that records of all selected farmland birds from semi-structured ornitho records tended to come from more densely populated areas than records from the structured monitoring scheme; median CBBS: 105 (Q25-Q75: 51.2 – 276) inhabitants per km^2^ whereas median semi-structured ornitho: 177 (Q25-Q75: 82 – 525) inhabitants per km^2^. The yearly distributions of human population density from the records were, however, constant over the analysed period (Appendix Fig. S11), in other words our estimation of the temporal trends is not biased due to a shift in the records towards more populated areas, potentially differing between the different datasets over time.

Another possibility of induced locational bias is the difference in the spatial coverage between structured monitoring schemes and online databases. For instance, as a crude comparison, 1,700 routes were sampled from the CBBS in Germany, while information from 25,000 and 184,000 “sites” (grid cells) were available from semi-structured and unstructured data. The larger spatial coverage in data arising from online databases could act as an “early detection” system, reporting negative and positive trends before these are reaching populations followed by the structured monitoring scheme (Altwegg and Nichols 2019)

The potential of semi- and unstructured data to act as an early detection system is usually impaired by the high skewness of the number of site visits: few interesting sites are repeatedly sampled by a lot of participants while the vast majority of sites have only one or a few records making it difficult to disentangle temporal changes and site effects (Isaac et al. 2014). One option to correct this skewness is to identify areas and habitats consistently undersampled from online databases and send trained observers to fill these data gaps in order to better allocate money and effort (Tulloch et al. 2013). To identify undersampled areas, a priori, the recording process from citizen scientists needs to be modelled and predicted (Johnston et al. 2020). Another option would rely on incentives such as rewards (monetary or not) or dynamic maps indicating undersampled areas to encourage volunteers to provide records in those areas or habitats (Xue et al. 2016). In this study we decided to heavily filter our data. The millions of available records were discretized to grid cells and only one record per grid cell, per species and per year was passed on to the models. The assumption being that this record represents the population status for that grid cell or site. By doing so, we heavily rely on the accuracy of that single observation, but at the same time we drastically down-weight the impact of the spatial sampling bias. Our data filtering approach aimed to eliminate possible migrants in the semi- and unstructured dataset across years by removing records outside of the species-specific breeding period. Species phenology is, however, affected by several environmental parameters, and breeding periods may change from year to year (Tøttrup et al. 2008).

In 2013, abundance indices derived from unstructured data showed large peaks in some species in March (Fig S12). This was probably due to bad weather conditions during migration resulting in large numbers of individuals resting in Germany. To account for this, we used an extra filter on top of the breeding period, the maximum number of breeding birds recorded in the structured monitoring scheme as a threshold of maximum bird abundance per grid cell in the semi-and unstructured dataset. This demonstrates the importance of carefully considering data filtering, the necessity of flexible filtering approaches and good knowledge of potential biases in semi- and unstructured data sources.

### Increased accuracy with Bayesian hierarchical models especially for less common species

The accuracy of the model, the difference between the observed breeding bird abundances and the model predicted values, was not affected by the integration of semi- and unstructured data. Rather we found that the TRIM models were usually less accurate than the Bayesian hierarchical models, especially for the less common species where Bayesian hierarchical models were up to 75% more accurate than TRIM. This implies that adding more information to estimate the yearly changes in bird abundances did not lead to better agreement between the model predicted values and the observed data but rather that fundamental differences in the fitting algorithm (Generalized Estimating Equations vs Hamiltonian Monte Carlo) can lead to differences in accuracy. This could be due to the fact that the fitted models were rather rigid, differences in breeding abundances could only come from site or from year effects. Adding complexity and flexibility to the model such as through site-level covariates and increasing the number of model parameters that are jointly estimated across several datasets could lead to increased predictive accuracy under data integration (Simmonds et al. 2020) but further evidence from real-life data is needed.

### Increased precision with data integration

Precision, the degree of uncertainty around the predicted temporal trends, increased with data integration probably due to larger sample sizes. Compared to models fitted with structured data only, the increase in precision was around +25% when semi-structured data only were integrated and +50% when both semi-structured and unstructured data were integrated. This pattern was not affected by species prevalence or body mass and only in one model by population size. More precise estimation of temporal trends increases the power to detect declines or increase in breeding bird population (Zipkin et al. 2017, Bowler et al. 2019). Bird population trends are widely used as indicators of the state of nature, and especially farmland birds’ trends have been used repeatedly to assess the sustainability of farming practices, and the efficacy of Agri-Environmental schemes and agricultural policies. (Gregory et al. 2005). In this context models integrating additional data from online databases to structured monitoring data could complement the current modelling approach to ensure better or earlier detection of positive or negative trends in bird populations. This could especially be important for species with low sample size in systematic monitoring schemes e.g. due to small populations and range sizes, or rapid population declines.

### About data quality

Working with semi- and unstructured data requires to consider certain key characteristics of these datasets either through data pre-processing (Johnston et al. 2019) or by using specific models (Isaac et al. 2014). Semi- and unstructured data come with different levels of information, purely opportunistic records where only the “what-when-where” are being recorded is difficult to analyse. Adding information on the “how”, especially on the effort that went into the collection of the records is critical for efficiently modelling these data (Johnston et al. 2019). In this regard the use of complete checklists in citizen science biodiversity records such as pioneered in eBird (Sullivan et al. 2014) is particularly useful and can provide reliable information on, for instance, intra- and inter-annual spatio-temporal trends (Kelling et al. 2019). Our results demonstrate that the integration of complete checklist or semi-structured data with structured data provides both reliable and more precise estimation of temporal trends in bird populations. The integration of unstructured data increased precision strongly but reliability of population trends compared to benchmark models varied strongly across species. Finally, data integration methods following joint likelihood approaches (Pacifici et al. 2017) can also include a weighting of the different dataset in order to account for changes in data quality (Renner et al. 2019). Future model development could explore such weighting functions to prevent purely opportunistic data to overwhelm the sparse but good quality structured dataset. Another option, also developed in the context of citizen science data, is spatio-temporal subsampling to account for differences in data amount across space and time (Fink et al. 2020).

## Conclusions

The potential of integrating citizen science data with data from structured, standardized sampling schemes has so far mainly focused on estimating spatial distribution of the species (Pacifici et al. 2017, Miller et al. 2019, Isaac et al. 2020). Yet, data integration can be performed to estimate any model components, and is commonly used for other types of ecological models such as in Integrated Population Models to estimate demographic parameters such as survival or reproduction from different data streams (Sun et al. 2019). Here, we demonstrated how integration of citizen science data can improve the precision of the predicted temporal trends in bird populations. Furthermore, models integrating semi-structured data predicted consistent trends in more than 75% of the studied species compared to models built on structured data only. We found that the greatest potential of data integration lies in models using the structured dataset together with semi-structured data given that the latter come with key information on the data collection process that is absent from unstructured records. Data integration approaches allow us to go a step further compared to previous studies comparing models fitted with structured vs. unstructured data (Kamp et al. 2014, Boersch-Supan et al. 2019). Importantly, parameter estimation in data-integrated models can leverage information across datasets while accounting for varying sampling process and observational errors (Isaac et al. 2020). In addition, the effect of more explanatory variables could be jointly assessed as the number of observation and the observed gradient in the co-variates increases. Here we explored data integration potential using farmland birds, a group of species that is relatively well covered in structured monitoring programs (Proença et al. 2017). It can be expected that data integration would yield even bigger benefit in taxa enjoying lower sampling attention and resulting scarce data availability from structured monitoring programs (Boersch-Supan et al. 2019).

## Data availability statement

The CBBS and ornitho data are available for research upon reasonable request. The observation and eBird data are freely available. The Rscripts to reproduce the fitted models and the main figures are available in the following repository: https://doi.org/10.5281/zenodo.4287172

